# An obligate aerobe adapts to hypoxia by hybridising fermentation with carbon storage

**DOI:** 10.1101/2023.09.11.557286

**Authors:** David L Gillett, Tess Hutchinson, Manasi Mudaliyar, Thomas D. Watts, Wei Wen Wong, Jake Locop, Luis Jimenez, Iresha Hanchapola, Han-Chung Lee, Erwin Tanuwidjaya, Joel R. Steele, Ralf B. Schittenhelm, Christopher K. Barlow, Rhys Grinter, Debnath Ghosal, Perran L. M. Cook, Chris Greening

**Affiliations:** Department of Microbiology, Monash University, Clayton, VIC, Australia; School of Chemistry, Monash University, Clayton, VIC, Australia; Monash Proteomics and Metabolomics Facility, Monash University, Clayton, VIC, Australia; Department of Biochemistry and Pharmacology, Bio21 Institute, The University of Melbourne, Parkville, VIC, Australia; ARC Centre for Cryo-electron Microscopy of Membrane Proteins, Bio21 Molecular Science and Biotechnology Institute, University of Melbourne, Parkville, VIC, Australia

## Abstract

In soil ecosystems, obligately aerobic bacteria survive oxygen deprivation (hypoxia) by entering non-replicative persistent states. Little is known about how these bacteria rewire their metabolism to stay viable in these states. The model obligate aerobe *Mycobacterium smegmatis* maintains redox homeostasis during hypoxia by mediating fermentative hydrogen production. However, the fate of organic carbon during fermentation, and the associated remodeling of carbon metabolism, is unresolved. Here we systematically profiled the metabolism of *M. smegmatis* during aerobic growth, hypoxic persistence, and the transition between these states. Using differential isotope labelling, and paired metabolomics and proteomics, we observed rerouting of central carbon metabolism through the pentose phosphate pathway and Entner-Doudoroff pathway during hypoxia. We show that *M. smegmatis* excretes high levels of hydrogen concurrently with upregulating triacylglyceride synthases and accumulating glycerides as carbon stores. Using electron cryotomography (cryo-ET), we observed the presence of large spheroid structures consistent with the appearance of lipid droplets. Thus, in contrast to obligately and facultative anaerobic fermentative bacteria, *M. smegmatis* stores rather than excretes organic carbon during hypoxia. This novel hybrid metabolism likely provides a competitive advantage in resource-variable environments by allowing *M. smegmatis* to simultaneously dispose excess reductant during hypoxia and maintain carbon stores to rapidly resume growth upon reoxygenation.

## Introduction

Most soil bacteria exist in non-replicating persistent states, also known as dormancy. In these states, bacteria do not expend energy on growth and replicate, but still require some minimal metabolic activity to sustain cellular integrity and maintenance. Limitation or variability for key resources, for example organic carbon, nitrogen, or oxygen, is the primary reason microbes transition from growing to dormant states^1–3^. In many soil environments, oxygen concentrations sharply vary across space and time, and influence the biological structure at both microscopic and macroscopic scales. Many soil bacteria are facultative anaerobes that adapt to oxygen limitation (hypoxia) by growing through anaerobic respiration or fermentation instead. However, the dominant bacteria in most surface soils are seemingly obligate aerobes that can only survive rather than grow during hypoxia^4^. For example, *Mycobacterium* is a cosmopolitan genus of obligate aerobes that comprise approximately 1% of soil bacteria and are notorious for their capacity to survive extended periods of hypoxia^4–6^. To date, relatively little is known about how mycobacteria and other aerobic soil bacteria adapt their metabolism to maintain viability under hypoxia.

Recent studies suggest soil mycobacteria depend on metabolic flexibility to adapt to variations in resource limitation^7–9^. We’ve shown that *M. smegmatis* switches between growth through aerobic respiration and persistence by fermentation in response to variations in oxygen availability. When oxygen is limited, this bacterium disposes of excess electrons by reducing protons to H_2_ using the fermentative group 3b [NiFe]-hydrogenase Hyh^7^. *M. smegmatis* is proposed to recycle H_2_ produced during hypoxia using the respiratory group 2a [NiFe]-hydrogenase Huc when electron acceptors become available. Production of both hydrogenases is activated under hypoxia by the oxygen- and redox-sensing DosS/T-DosR two-component system^7^. Deletion of both respiratory and fermentative hydrogenases disrupts redox homeostasis and decreases survival^7,10^. While this study provided the first report of fermentation in an obligate aerobe, it did not address the fate of the carbon end-products. Most previous studies have investigated fermentation in the context of growth rather than persistence^11–13^; for example, in the well-known examples of ethanol and lactic acid fermentation, the fermentative end-product is secreted outside of the cell, preventing intracellular accumulation of organic waste molecules^12,13^. However, it is unclear whether *M. smegmatis* also excretes end-products or instead stores them. Organic carbon is still potentially available for anabolic metabolism after it has been partially oxidised to drive ATP production. Moreover, anabolic storage pathways also serve as a sink of reductant that can be used to regenerate electron carriers, but it is difficult for cells to maintain redox balance through this process alone. In this manner, fermentative hydrogenases may serve as a redox valve by allowing the cell to release excess reductant and regenerate redox cofactors as required.

The synthesis and accumulation of triacylglycerols (TAGs) is an important part of mycobacterial persistence^14–19^. TAGs are composed of three fatty acid chains joined by ester linkages to a glycerol backbone; a lack of any hydrophilic moiety makes them exceedingly hydrophobic so that they readily form droplets, and the highly reduced fatty acid chains can be readily oxidised to extract energy. These properties make TAGs an ideal cellular store of energy and carbon, as lipid droplets are self-compartmentalising and extremely energy dense. Lipid droplets, also referred to as intracellular inclusions, were first reported in mycobacteria more than 70 years ago and have since been consistently observed in mycobacteria via light and electron microscopy^15,20–24^. These droplets are strongly associated with stress responses and persistence in mycobacteria^15,18,25^ and TAGs have been identified as their dominant constituent^18,26,27^. Mycobacterial genomes encode multiple TAG synthases (Tgs), which catalyse the final step of TAG synthesis (the acylation of diacylglycerols) and these are among the most upregulated genes in mycobacteria during hypoxia, with Tgs1 the most active of the synthases part of the regulon of the hypoxic response regulator DosR in both *M. tuberculosis* and *M. smegmatis* ^7,28–30^. As well as a storage molecule, TAGs are also abundant within and on the surface of the mycobacterial cell envelope^31,32^, as are TAG synthases and TAG export proteins^17,32,33^ and their hydrophobicity is thought to contribute to the highly impermeable nature of the mycomembrane. Our understanding of TAG metabolism in mycobacteria predominately arises from the study of *M. tuberculosis,* which builds substantial stores of TAGs during infection, via both biosynthesis and the import of TAGs and fatty acids from host lipid-loaded macrophages^15–17,19^. In contrast, the relationship between TAG metabolism and the persistence of environmental mycobacteria is poorly studied.

Building on these previous studies, we hypothesised that these processes are coupled: *M. smegmatis* may adapt to hypoxia by hybridising hydrogenogenic fermentation with TAG synthesis. We systematically tested this by measuring carbon degradation pathways and hydrogen production, conducting paired metabolomics and proteomics, and visualising lipid droplets in cells before, during, and following the transition to hypoxia. Together, we provide strong evidence that *M. smegmatis* partially oxidises organic carbon during hypoxia by disposing excess reductant as hydrogen and storing organic carbon primarily as TAGs. We propose that this novel hybrid mode of metabolism is ideally suited for bacteria adapting to frequent oxic-hypoxic transitions.

## Results

### *M. smegmatis* partially oxidises carbon during fermentative H_2_ production

*M. smegmatis* produces H_2_ during oxygen depletion using the fermentative hydrogenase Hyh^7^, though the timing of this process and the extent of H_2_ accumulation have not been established. To understand the dynamics of hydrogen production during the transition to hypoxia, we cultured *M. smegmatis* in Hartman’s de Bont (HdeB) minimal media with 0.4% w/v glucose in sealed serum vials with ambient air and measured the concentration of H_2_ and O_2_ in the headspace as O_2_ was gradually depleted (**Fig. 1A**). OD_max_ was reached when headspace O_2_ concentration reached ∼2%, and O_2_ was gradually consumed to below the detection limit (∼0.10%) over the next 24 hours, indicating transition to hypoxia-induced dormancy. This is broadly consistent with existing experimental models of hypoxia-induced mycobacterial dormancy^34,35^. The onset of net hydrogen production coincided with the consumption of O_2_ below detection limits. H_2_ concentration steadily rose over the next 48 hours before plateauing between 500-1000 ppmv, indicating that organic carbon catabolism is sustained through the transition to hypoxia and is supported by fermentative H_2_ production once respiratory electron acceptors are exhausted (**Fig. 1A**). The concentration of accumulated H_2_ is a magnitude higher than previously reported using similar methodology^7^, and suggests that fermentative H_2_ production is a major process during hypoxia in *M. smegmatis*.

**Figure 1:**
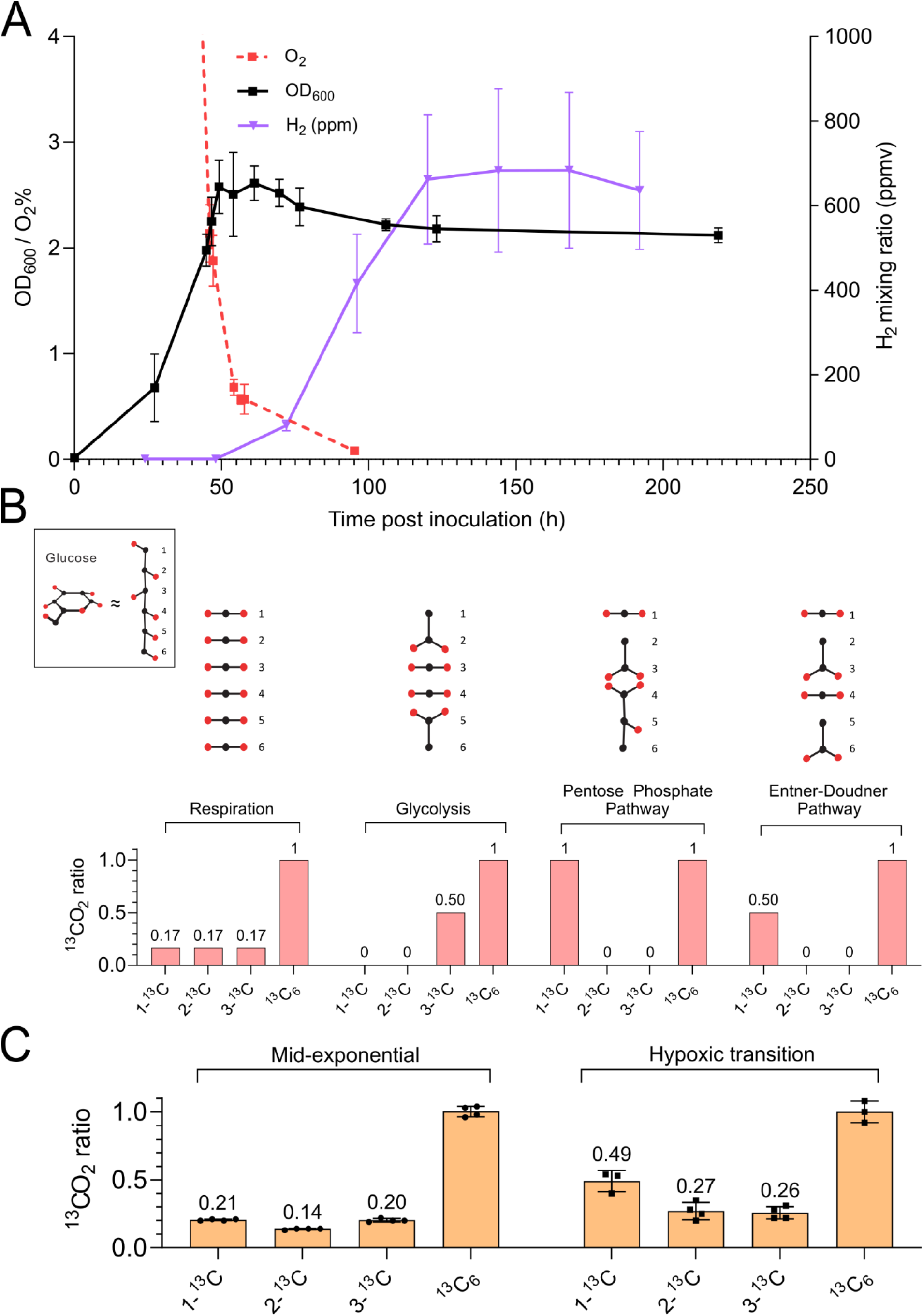
*M. smegmatis* fermentatively produces hydrogen and partially oxidises glucose via the pentose phosphate pathway during oxygen depletion. (A) Density (OD_600_), O_2_ and H_2_ of *M. smegmatis* cultures (n=4) during gradual oxygen depletion were monitored. Density and O_2_ share the same axis and range. Growth arrest (OD_max_) is reached at [O_2_] = ∼2% while H_2_ accumulation coincides with onset of anoxia at ∼70 hours post inoculation. Error bars represent standard deviation from the mean. (B) Catabolism of glucose through either glycolysis, the Pentose Phosphate Pathway (PPP) or the Entner-Doudoroff Pathway (EDP) results in carbons at different positions being oxidised to CO_2_. Treatment with differentially isotope labelled glucose and comparing the ratios of ^13^CO_2_/^12^CO_2_ by using isotope ratio mass spectrometry (IRMS) indicates contributions of different catabolic pathways to CO_2_ production. (C) *M. smegmatis* (n=4) was treated with differentially ^13^C-labelled glucose at mid-exponential phase (OD_600_ ∼0.8) and the hypoxic transition (late-exponential phase; OD_600_ ∼2.0).

In contrast to respiration where organic carbon is completely oxidised to CO_2_, during fermentation organic carbon is incompletely oxidised (**Fig. 1B**). To determine if the degree of oxidation of organic carbon is consistent with fermentation during the transition to hypoxia-induced dormancy, we incubated *M. smegmatis* cultures with 1-^13^C-glucose, 2-^13^C-glucose, 3-^13^C-glucose and ^13^C_6_-glucose for 5 hours during exponential growth, stationary phase, and during the onset of hypoxia (OD_600_= ∼2.5), and then measured the ratio of ^13^CO_2_:^12^CO_2_ using isotope ratio mass spectrometry (IRMS)^36^. If all six carbons of glucose are oxidised, as during respiration, each carbon will contribute 1/6^th^ of the total ^13^CO_2_ signal produced by ^13^C_6_-glucose, while incomplete oxidation will result in individual carbon positions contributing disproportionately to the total ^13^CO_2_ signal (**Fig. 1B**). During exponential growth, the proportion of signal attributable to 1-^13^C, 2-^13^C and 3-^13^C was 0.21, 0.14 and 0.20 of the total ^13^C_6_ signal respectively, indicating that aerobic respiration was the dominant catabolic process, as expected, while during the hypoxic transition there was a disproportionate increase in the ^13^CO_2_ signal attributable to 1-^13^C glucose (0.49 of ^13^C-6) (**Fig. 1C**). This is consistent with glucose being incompletely oxidised as it passes through either the pentose phosphate pathway (PPP) or the Entner-Doudoroff pathway (EDP), two alternative pathways for the oxidation of glucose to pyruvate (**Fig. 1B**). Substantial ^13^CO_2_ signal was still attributable to 2-^13^C and 3-^13^C, indicating that respiration was also occurring, which is unsurprising considering O_2_ is still present before OD_max_.

We also treated hypoxic mid-stationary phase *M. smegmatis* cultures (3 days post OD_max_) with differentially labelled 13-C glucose but detected no ^13^CO_2_ signal above background, even after an extended incubation of 5 days. This indicates that *M. smegmatis* stopped consuming exogenous glucose in the media after prolonged anoxia, possibly because of reduced metabolic activity or a switch to the catabolism of internal carbon stores.

### Early proteome remodelling drives a sustained shift in metabolism during hypoxia

To gain further insights into the fate of partially oxidised carbon during fermentation, and associated changes to broader metabolism, we conducted paired comparative metabolomics and proteomics on lysates from *M. smegmatis* cultures harvested during aerated exponential growth, the transition to hypoxia (0.5% O_2_), and sustained hypoxia (mid-stationary phase, 3 days post OD_max_). We observed large changes across all comparisons: 397 metabolites and 310 proteins were differentially abundant between hypoxic transition vs exponential phase, 610 metabolites and 307 proteins between stationary phase vs exponential phase and 429 metabolites and just two proteins between stationary phase vs hypoxic transition (**Fig. 2A and 2C**). The changes in the metabolome were consistently larger than those for the proteome, and with just two proteins reaching the cut-off criteria (>2 fold change, P < 0.01), virtually no changes to the proteome were observed between the hypoxic transition and stationary phase. This indicated that proteome remodelling is completed early in the response of *M. smegmatis* to oxygen depletion, but drives a prolonged and sustained shift in metabolism. This is somewhat unexpected, as large transcriptional changes are sustained throughout the response of *M. tuberculosis* to oxygen depletion^28,37^.

**Figure 2:**
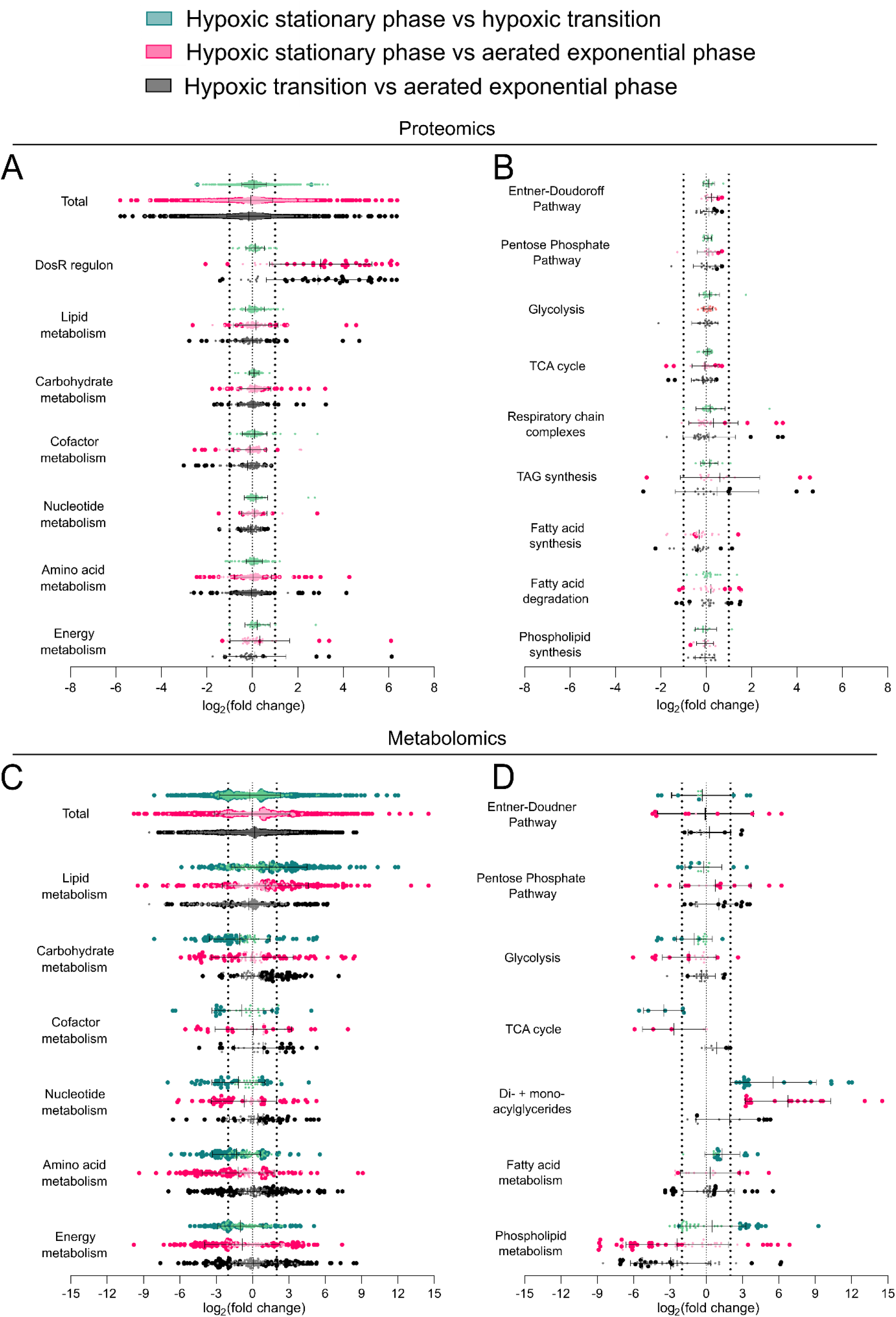
Comparative proteomic (top panels) and metabolomic (bottom panels) analysis of *M. smegmatis* cultures (n=4) during mid-exponential phase (EXP), hypoxic transition (TR) and hypoxic stationary phase (ST). Comparisons are separated into TR vs EXP (black), ST vs EXP (dark pink) and ST vs TR (dark green). Proteins and metabolites with statistically significant abundance across comparisons (proteomics, p<0.05 (A-B); metabolomics, p < 0.01 (C-D)) are represented with large dark coloured dots. Proteins and metabolites that were detected but were not statistically different are represented with smaller dots in lighter shades. The fold change (FC) threshold is represented with thick dotted lines (proteomics, FC > 1 (log_2_) (A-B); metabolomics, FC > 2 (log_2_). All error bars represent standard deviation from the mean. Differentially abundant proteins are separated into predicted metabolic functional groups based on KEGG pathways annotations (A) and KEGG module annotations (B). Differentially abundant metabolites are separated into predicted metabolic function based on IDEOM annotations (C) and KEGG modules annotations (D). Predicted annotations are superseded with experiment-based annotations where available.

Consistent with the broader literature, the most highly upregulated proteins during oxygen deprivation were members of the DosR regulon (**Fig. 2A**), with the most upregulated proteins being HspX and the FAD-sequestering protein Fsq^38^. The α, β and δ subunits of Hyh were all upregulated between 2.8-fold and 38-fold during the hypoxic transition (hypoxic transition vs exponential phase), consistent with our finding that net hydrogen production is observed shortly after oxygen is depleted below the detection threshold. Two DosR-regulated triacylglycerol synthases that drive TAG synthesis in *M. tuberculosis* during hypoxia^29^ were upregulated 16- to 26-fold during hypoxia. The α and β subunits of the uptake hydrogenase Huc were mildly upregulated (∼2.5 fold), which is consistent with previous studies that hydrogen produced by Hyh can be recycled by Huc if a terminal electron acceptor becomes available^7^.

For each comparison, we broadly categorised metabolites and proteins by predicted function according to KEGG and IDEOM pathway annotations (**Fig. 2A and 2C**). For metabolites, we observed large changes in all comparisons for every category examined. For energy metabolism, amino acid metabolism, cofactor lipid metabolism, and carbohydrate metabolism, multiple metabolites increased and decreased in abundance by at least 8-fold. We detected an overall increase in lipid metabolism in the stationary vs exponential phase comparison (mean of 2.1-fold) and carbohydrate and cofactor metabolism in the hypoxic transition vs exponential phase comparison (means of 2.1- and 2.0-fold respectively) and an overall decrease in nucleotide metabolism, amino acid metabolism and peptides in the stationary phase vs exponential phase comparison (means of -1.5, -2.1 and -1.6 fold). Changes in the proteome to metabolic pathways during oxygen depletion were generally much milder than for the metabolomes, and except for the strong upregulation of the DosR regulon, we discerned no pattern in the comparisons (**Fig. 2 and Fig. 3**).

**Figure 3:**
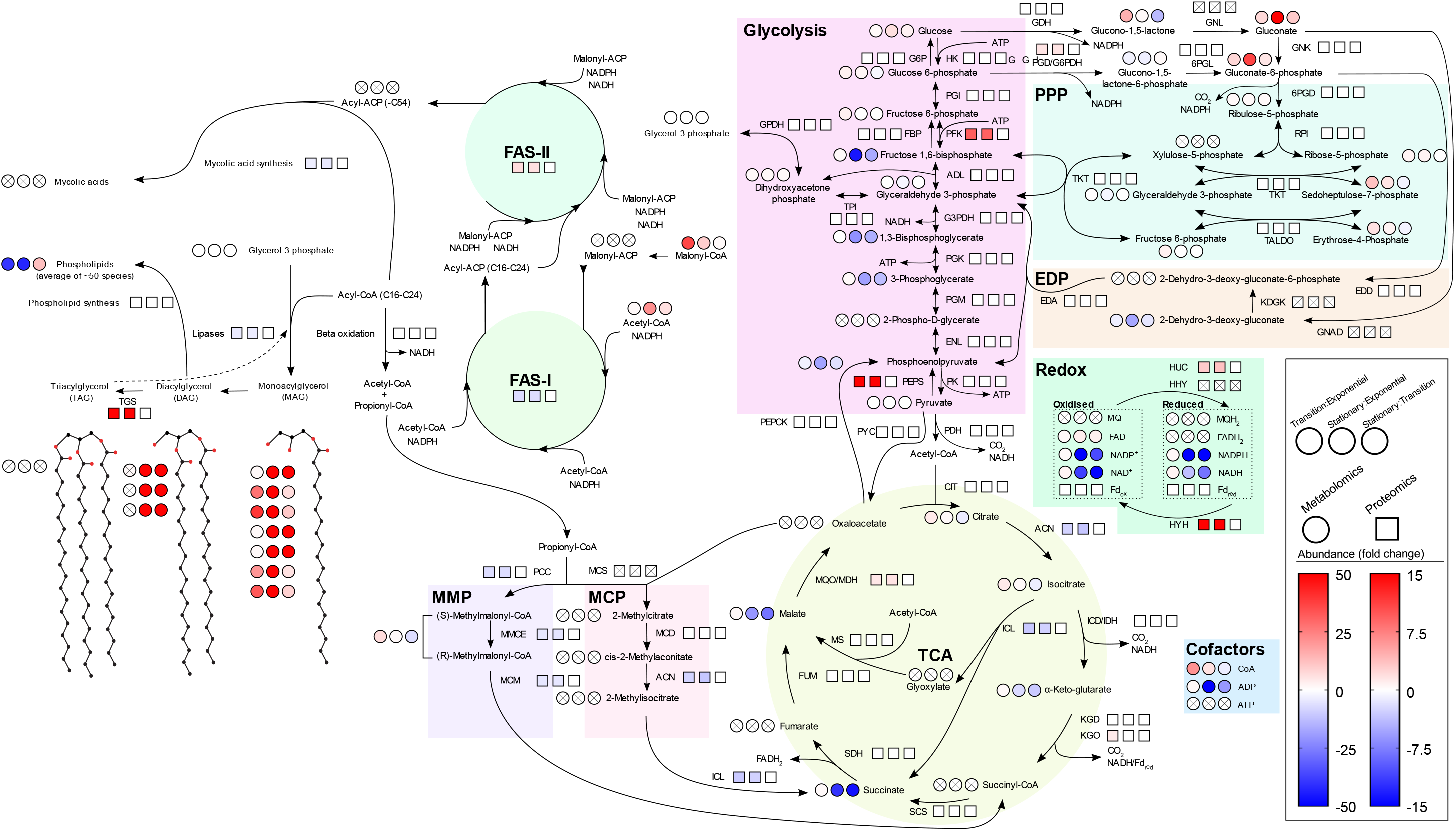
Summary of metabolic changes during oxygen depletion in *M. smegmatis*, as determined by comparative proteomics and metabolomics (Fig. 2). As described in legend (bottom right), heat map displayed as circles (metabolites) and squares (proteins) represents fold change in abundance for features meetings their respective cut-off threshold (proteomics, P < 0.05, FC > 2; metabolomics, p < 0.01, FC > 4). Features not identified in experiment are marked with X and white background. For clarity, oxidised cofactors (e.g., NAD(P)^+^) and ADP are not included. Abbreviations: ACN: aconitase; ACP: acyl-carrier protein; ADL: fructose-bisphosphate aldolase; ADP: adenosine diphosphate; ATP: adenosine triphosphate; CIT: citrate synthase; CoA: Coenzyme A; EDA: 2-dehydro-3-deoxyphosphogluconate aldolase; EDD: phosphogluconate dehydratase; EDP: Entner-Doudoroff pathway; ENL: enolase; FAD: flavin adenine dinucleotide; FAS: fatty acid synthase; Fd_red/ox_: ferredoxin (reduced / oxidised) FGD: F_420_-dependent G6PDH; FUM: fumarase; GAPDH: glyceraldehyde phosphate dehydrogenase; G6PDH: glucose-6-phosphate dehydrogenase; GDH: glucose-1 dehydrogenase; GNK: gluconokinase; GNL: gluconolactonase; GPDH: glycerol-3-phosphate dehydrogenase; HHY: respiratory group 1h [NiFe]-hydrogenase HK: hexokinase; HUC: respiratory group 2a [NiFe]-hydrogenase; HYH: fermentative group 3b [NiFe]-hydrogenase; ICD/IDH: isocitrate dehydrogenase; ICL: isocitrate lyase; KDGK: 2-dehydro-3-deoxygluconokinase; KDH: ketoglutarate dehydrogenase; PGK: phosphoglycerate kinase; PGM: phosphoglycerate mutase; PK: pyruvate kinase; MCS: methylcitrate synthase; MCD: methylcitrate dehydratase; MCM: methylmalonyl-CoA mutase; MCP: methylcitrate pathway; MDH: malate dehydrogenase; MMCE: methylmalonyl-CoA epimerase; MMP: methylmalonyl pathway; MS: malate synthase; NAD(H): nicotinamide adenine dinucleotide; NADP(H): nicotinamide adenine dinucleotide phosphate: PCC: propionyl-CoA carboxylases; PDH: pyruvate dehydrogenase; PEPCK: phosphoenolpyruvate carboxykinase; PEPS: phosphoenolpyruvate synthase PFK: phosphofructokinase; PGI: phosphoglucoisomerase; PPP: Pentose Phosphate Pathway; PYC: pyruvate carboxylase; RPI: ribulose-5-phosphate isomerase; SCS: succinyl-CoA synthetase; SDH: succinic dehydrogenase; TALDO: transaldolase; TCA: tricarboxylic acid cycle; TKT: transketolase; TGS: triacylglycerol synthase; TPI: triosephosphate isomerase; 6PGD: 6-phosphogluconate dehydrogenase;

### *M. smegmatis* drives TAG synthesis during hypoxia

To investigate changes to metabolic pathways relevant to fermentation and TAG synthesis, we examined changes to central carbon metabolism and fatty acid metabolism. Large accumulations of monoacylglycerols (MAGs) and diacylglycerols (DAGs) occurred during the hypoxic transition and hypoxic stationary phase (**Fig. 2B**). This is consistent with the strong upregulation of two TAG synthases, though it may also reflect the biosynthesis or degradation of other glycerolipids such membrane phospholipids. We observed a large decrease in phospholipids, indicating that accumulation of MAGs and DAGs is not due to an upregulation of phospholipid biosynthesis, and suggesting that phospholipid degradation could be a major source of the MAGs and DAGs. While we detected no TAGs, this reflects their highly hydrophobic nature is incompatible with our extraction and LC-MS protocol.

Intermediates of fatty acid metabolism were overall more abundant during oxygen depletion, with several free fatty acid species increasing in abundance by >20 fold (26:0, 19:1, 17:0, 14:0 and 16:1), indicating that either glycerolipids were being degraded to provide free fatty acids, or new fatty acids were being synthesised *de novo*. Malonyl-CoA, the precursor of malonyl-ACP which is used by fatty acid synthase I (FAS I) to initiate fatty acid chain synthesis (C16-C24) and by fatty acid synthase II (FAS II) to produce long chain fatty acids (up to C56) as precursors for mycolic acids, major components of the mycobacterial cell wall^39^, increased in abundance by 36-fold in hypoxic transition and 10-fold in hypoxic stationary phase compared to exponential phase. We observed a mild decrease in proteins involved in mycolic acid synthesis during oxygen depletion (<2-fold), but did not identify any mycolic acid synthesis intermediates, although this is likely be due to their highly hydrophobic nature preventing mass spectrometric detection. Overall, the accumulation of MAGs and DAGs is likely driven by a combination of both de novo fatty acid synthesis and the degradation of phospholipids. Although we cannot exclude the possibility of increased mycolic acid synthesis, the metabolomic and proteomic changes are overall consistent with a dramatic upregulation of TAG biosynthesis during hypoxia concurrent with fermentative H_2_ production.

### Rewiring of central carbon metabolism and reduction of cofactor levels during hypoxia

Changes to central carbon metabolism during hypoxia were characterised by an overall decrease in abundance of intermediates of glycolysis and the TCA cycle, and an increase in intermediates of the PPP and EDP, consistent with our differential isotope experiment. Interestingly, the abundance of glycolysis and TCA cycle intermediates only decreased in the hypoxic stationary phase vs exponential phase comparison, whereas the abundance of PPP and EDP intermediates increases in the hypoxic transition vs exponential phase (**Fig. 3**). The most dramatic changes were observed in gluconate and 6-phosphogluconate, precursors of both the PPP and EDP, both of which were markedly more abundant during hypoxic transition (7.4- and 7.7-fold, respectively) and hypoxic stationary phase phases (74-fold and 37-fold, respectively) (**Fig. 3**). Compared to exponential phase, PPP-intermediates sedoheptulose 7-phosphate and erthryose 4-phosphate were more abundant during the hypoxic transition (12-fold and 5.6-fold, respectively) and hypoxic stationary phase (5-fold and 2.2-fold, respectively), while the EDP intermediate 2-dehydro-3-deoxy-gluconate was less abundant during the hypoxic transition (3.5-fold) and hypoxic stationary phase (17-fold). This suggests increased flux through the PPP, rather than the EDP. We were unable to identify the key EDP intermediate 2-dehydro-3-deoxy-gluconate-6-phosphate (**Fig. 3**).

We saw a marked accumulation of gluconate, gluconolactone and gluconate-6-phosphate during oxygen depletion, early intermediates in both the EDP and PPP. Interestingly, the accumulation of gluconate and the transient accumulation of gluconolactone suggests an active gluconate shunt during oxygen depletion, which circumvents the first committed step of the PPP and ED: the oxidation of glucose-6-phosphate to glucono-1,5-lactone-6-phosphate by the glucose-6-phosphate dehydrogenase (G6PDH)^40^ (**Fig. 3**). The gluconate shunt is poorly studied in bacteria, and while *M. smegmatis* has genes predicted to encode the enzymes of the gluconate shunt (MSMEG_0655: glucose 1-dehydrogenase (GDH); MSMEG_1274 and MSMEG_2561: gluconolactonase (GNL); MSMEG_0453: gluconokinase), they have not been biochemically characterised. *M. smegmatis* encodes both a canonical NADP-dependent and a F_420_-dependent G6PDH, both of which were mildly upregulated (1.4- to 1.6-fold) during hypoxia, suggesting that the conventional oxidative PPP was active during hypoxia in addition to the gluconate shunt. Regardless, both routes produce reduced cofactors (i.e., NADPH,or F_420_H_2_) that supply reducing equivalents for anabolism, which is consistent with metabolic remodelling to support large-scale TAG synthesis. Furthermore, the accumulation of gluconate and gluconate-6-phosphate may reflect that metabolic flux is primarily rerouted through the PPP / EDP to produce reducing equivalents to support TAG synthesis, rather than intermediates for nucleotide and amino acid synthesis, for which we observed either no change or a decrease in abundance during oxygen depletion.

Based on the proteome, there were minimal changes to central carbon metabolism during oxygen depletion, with most enzymes exhibiting either no change or less than 2-fold change in abundance (**Fig. 2B and Fig. 3**). The exceptions to this were two products of the DosR regulon, MSMEG_3934 (upregulated 70-fold and 69-fold, hypoxic transition and hypoxic stationary phase, vs exponential phase, respectively), which encodes a putative phosphoenolpyruvate synthase (PEPS), and MSMEG_3947 (upregulated 9-fold for both hypoxic transition and hypoxic stationary phase, vs exponential phase), which encodes the reversible phosphofructokinase PfkB (**Fig. 3**). We also observed a 3.2- and 2.7-fold down regulation of isocitrate lyase (*icl)*, a key enzyme of the glyoxylate shunt and methylcitrate pathways (**Fig. 3**), in contrast to studies in *M. tuberculosis*. The potential roles of these enzymes are further discussed in **Supplementary Note 1**. We also observed a 4- to 8-fold upregulation of MSMEG_4710-4712, homologs of the subunits of the branched-chain keto acid dehydrogenase (BCKADH) complex for valine, leucine, and isoleucine degradation, consistent with the 5-fold increase in abundance of products ketoleucine and ketovaline during the hypoxic transition vs exponential phase comparison. Beyond these changes, the proteomes did not reflect the more dramatic changes in metabolite abundance. This potentially reflects that regulation of central carbon metabolism is largely post-transcriptional, for example through allosteric regulation, for example with studies in *M. tuberculosis* showing that PPP intermediates inhibit pyruvate kinase (PK) to further increase flux towards the PPP ^41^.

### *M. smegmatis* produces large droplets resembling lipid inclusions during hypoxia

To investigate if TAGs accumulated during hypoxia in *M. smegmatis* either within the cytoplasm or cell envelope, we used cryo-ET to investigate changes to the ultrastructure of *M. smegmatis* during oxygen depletion. We harvested *M. smegmatis* cultures at exponential phase (OD_600_ = ∼0.5, O_2_ = ∼18%), the hypoxic transition (OD_600_ = 2-2.5, O_2_ = 0.5 – 1%) and hypoxic stationary phase (5 days post OD_max_). During hypoxic stationary phase we consistently observed large and electron dense spheroid structures primarily clustered towards the bacterial cell poles, which are consistent with appearance of intracellular lipid inclusions in previous studies^18,26,27^ (**Fig. 4AB**). In contrast, during exponential phase, the cytoplasm of *M. smegmatis* cells were homogenous, except for density we attributed to ribosomes clustered towards the cytoplasmic membrane and the nucleoid (**Fig. 4CD**). We were unable to obtain tomograms of cells at the hypoxic transition. The distance between the inner membrane (IM) and outer membrane (OM) did not change substantially between exponential phase (∼24 nm) and hypoxic stationary phase (∼26 nm). This is consistent with the accumulation of DAGs and MAGs due to the biosynthesis and accumulation of TAGs, rather than the large-scale deposition of TAGs or other glycerolipids into the cell envelope.

**Figure 4:**
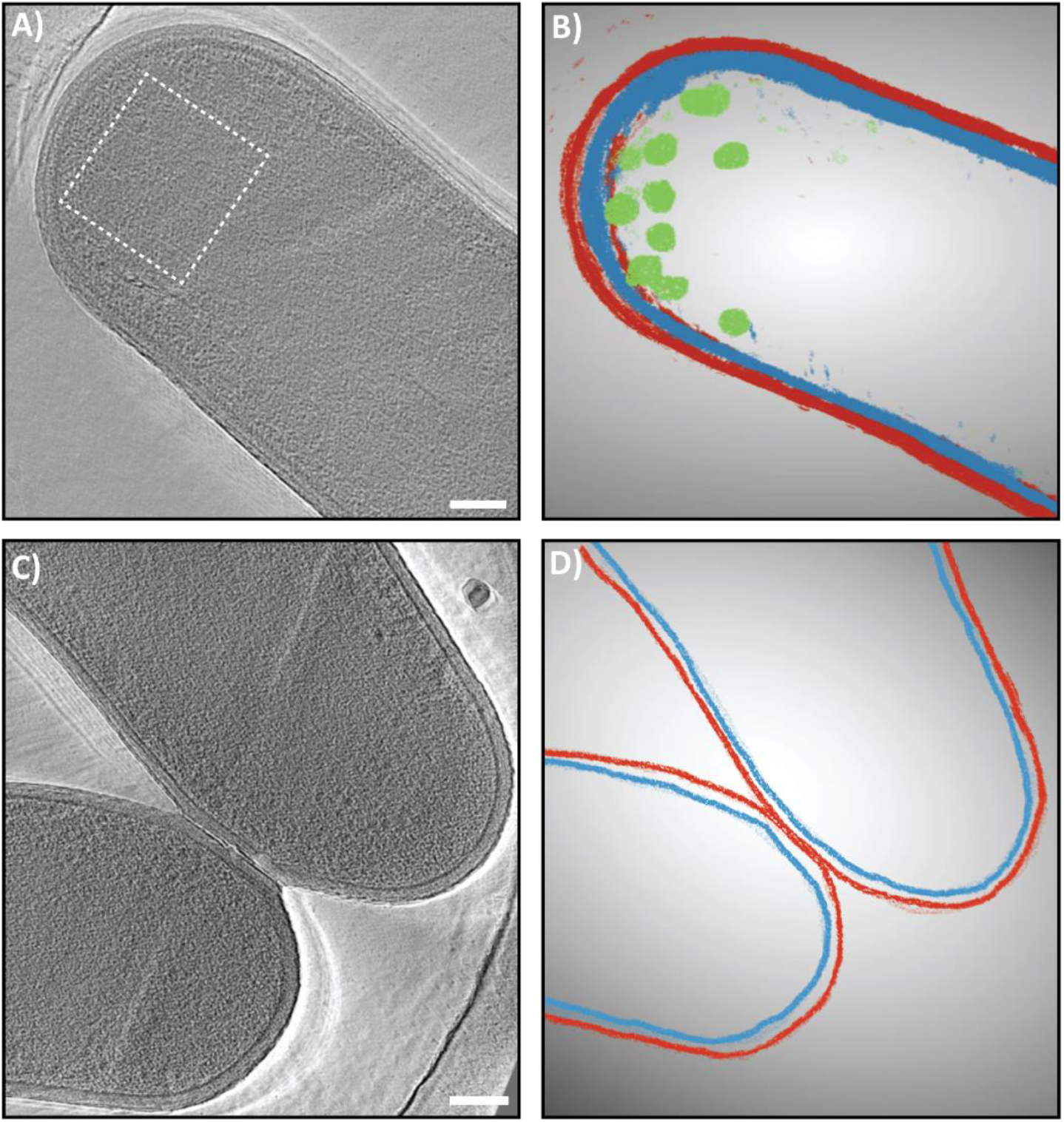
Cryo-ET of hypoxic and aerated *M. smegmatis* cultures. (A) Representative tomographic slice of hypoxic stationary phase (3 days post OD_max_) culture of *M. smegmatis*. White box indicates a region with several lipid droplets. (B) Segmentation analysis of the same tomogram showing lipid droplets (green), inner membrane (blue), outer membrane (red). (C) Representative tomographic slice of aerated mid-exponential phase culture of *M. smegmatis*, showing homogenous cytoplasm. (D) Segmentation analysis of the same tomogram shows inner membrane (blue) and outer membrane (red) but no lipid droplets. Scale bars 100 nm.

## Discussion

In this study, we characterised the metabolic remodelling associated with the transition of *M. smegmatis* to hypoxia. When exposed to a gradual onset of hypoxia, we found that *M. smegmatis* remodelled its proteome extensively during hypoxia but before oxygen was fully depleted (∼0.5%), and ceased growth between 2-3% oxygen, indicating that this remaining oxygen is used to support the transition to dormancy. Consistent with previous studies^7,30,34^, the most upregulated proteins during hypoxia were members of the DosR regulon. This included the fermentative hydrogenase Hyh. Consistently, we observed substantial accumulations of H_2_ in *M. smegmatis* cultures, with net H_2_ production beginning almost immediately after the depletion of oxygen below detection limits. This indicates that Hyh is highly active during oxygen depletion and could support the bulk of cofactor regeneration during ATP production through substrate-level phosphorylation. It should be noted that H_2_ accumulation is only observed after the complete depletion of oxygen. Previous studies show that Hyh is also active during early hypoxia, but the uptake hydrogenase Huc immediately recycles the H_2_, preventing its accumulation, indicating that *M. smegmatis* balances both aerobic respiration and fermentative H_2_ production to maintain redox homeostasis^7^. This agrees with our differential isotope labelling experiment, which indicated respiration and fermentation are both active during the early stages of hypoxia. The expression of Hyh during hypoxia could also be a proactive response to ensure fermentation begins as soon as all oxygen is depleted, therefore minimising any disruption to central carbon metabolism.

Our study strongly supports our hypothesis that *M. smegmatis* directs carbon partially oxidised during fermentative H_2_ production towards large reservoirs of TAGs during oxygen depletion. This is due to the observed accumulation of MAGs and DAGs, the strong upregulation of TAG synthases, the appearance of intracellular lipid inclusions during oxygen depletion, and the rerouting of carbon through the PPP and EDP to provide NADPH for fatty acid synthesis. These findings are broadly consistent with the accumulation of TAGs in *M. tuberculosis* during hypoxia^28^. The metabolic scheme shown in **Figure 3** shows how hypoxic cells rewire their carbon metabolism to dispose excess reductant as H_2_ and organic carbon as TAGs. Crucially, while TAG synthesis serves as a large sink of reducing equivalents for *M. smegmatis* during hypoxia, the simultaneous activity of hydrogen fermentation indicates that there is still an excess of reducing equivalents as the cell builds TAG stores, likely because the cell is only able to generate ATP with pathways that also produce NADPH or NADH, i.e., the PPP, EDP or glycolysis. To our knowledge, this is the first report of this hybrid mode of metabolism, though it is likely that other obligate aerobes adopt comparable strategies to adapt to resource variability.

The rapid and complex response to hypoxia, including hydrogen cycling and carbon storage, likely reflects the dynamic aeration conditions that obligate aerobes like *M. smegmatis* are exposed to. Soil constitutes a complex and irregular physical matrix with uneven distributions of oxygen and water at the microscopic level, and bacteria may be frequently exposed to oxygen depletion within their microenvironment^42–44^. Alongside genes encoding for aerobic respiration, fermentative [NiFe]-hydrogenase genes are prevalent within wetland, forest and grassland soils microbial communities, suggesting that fermentative H_2_ production could be a widespread strategy that aerobic bacteria use in response to oxygen depletion^4^. Moreover, in environments that are transiently rather than persistently deoxygenated, the production of reduced carbon stores rather than excretion of fermentative endproducts (e.g. volatile fatty acids) is likely to be adaptive for multiple reasons, namely: it provides a mechanism to acquire organic compounds from the environment even when oxygen is limiting; it enables rapid resumption of metabolism and growth upon reoxygenation; and it avoids release of organic compounds that may provide competitors (including facultative anaerobes) with an advantage. It is likely that these observations also extend to other ecosystems. For example, in coastal permeable sediments (i.e. sand) which frequently experience dramatic changes to redox conditions over small spatial scales^45^, fermentative H_2_ production and carbon storage appear to be biogeochemically and ecologically significant processes^46,47^. Building on these findings, future studies should further investigate the roles of fermentation and carbon storage in the context of dynamically aerated environments and microbial persistence.

## Material and Methods

### Culturing of *M. smegmatis*

*M. smegmatis* mc^2^155 was grown from frozen glycerol stocks on lysogeny broth agar plates supplemented with 0.05% (w/v) Tween80 (LBT) for 3-4 days at 37 C°. Starter cultures were inoculated with colonies from LBT plates and were grown in LBT media at 37 C° and shaking at 200 rpm overnight. Minimal media cultures were inoculated from turbid starter cultures to OD_600_ = 0.003 - 0.015 and grown in 30 mL Hartman’s de Bont (HdB) minimal media supplemented with 0.4% (w/v) glucose and 0.05% tyloxapol. Minimal media cultures and incubated at 37 °C with shaking at 150 rpm in 120 mL serums vials (Sigma, #33111-U) sealed with rubber stoppers (Sigma, #Z166065) and aluminium caps (Sigma, #Z114146)^7^.

### Measurement of hydrogen and oxygen concentrations during oxygen depletion

Measurements of hydrogen, oxygen and OD_600_ were taken from sealed cultures of *M. smegmatis* mc^2^155 from early stationary phase (∼24 hours post inoculation, OD600 = ∼0.4) to mid-stationary phase (3 days post OD_max_, OD_600_ = 2.5 - 3). Separate sets of replicate cultures were used for each type of measurement. To measure headspace O_2_ concentrations, a Retractable Needle-Type Oxygen Minisensor (PyroScience, #OXR430) was used with a FireSting Optical Oxygen meter (PyroScience, # FSO2-C4). The oxygen minisensor was calibrated using 12 mL exetainers purged with N_2_ and ambient air in sealed 120 mL serums vials incubated at 37°C. To measure headspace H_2_ concentrations, 200 µL of headspace was sampled using a gas-tight Luer lock 0.25 mL syringes (PhaseSep, # 050051-LL) flushed with N_2_ and immediately diluted via injection into a N_2_-purged 3 mL exetainer. To prevent contamination of oxygen depleted cultures with oxygen, sampling was conducted while culture vials were submerged in sterile milliQ water, Headspace H_2_ concentration was measured by injecting 2 mL of diluted headspace into a pulsed discharge helium ionisation detector (model TGA-6791-W-4U-2, Valco Instruments Company Inc.). A three-point calibration (100, 10 and 0 ppmv) curve was constructed and used to interpolate H_2_ concentrations.

### Differential isotope labelling

Sealed cultures of *M. smegmatis* were grown to mid exponential (OD_600_ = ∼0.7), hypoxic transition (OD_600_ = ∼2.5, approx. 5 hours pre OD_max_) and hypoxic stationary phase (3 days post OD_max_) in quadruplicate for each timepoint and condition. At each timepoint, cultures were treated with 50 µM of either 1-^13^C-, 2-^13^C-, 3-^13^C- or ^13^C_6_-glucose. To prevent contamination of oxygen depleted cultures with oxygen, sampling was conducted while culture vials were submerged in sterile milliQ water, and solutions of isotope labelled glucose and milliQ H_2_O were degassed via purging with N_2_ for at least 30 minutes in 12 mL gastight exetainers (Labco Exetainer) and treated using a gas-tight Luer lock 0.25 mL syringes (PhaseSep, # 050051-LL) flushed with N_2_. Exponential and hypoxic transition cultures were incubated for 5 hours post-treatment, and mid-stationary phase cultures were incubated for 5 days. Immediately after incubation, 12 mL of each culture was transferred to a gastight glass vial (12 mL, Labco Exetainer) with no headspace and immediately terminated via treatment with 20 µL 6% HgCl_2_ solution. Prior to ^13^CO_2_ analysis, 4 mL of sample in the 12 mL vial was replaced with helium. Phosphoric acid (12.5 mM) was added to the sample to convert dissolved inorganic carbon to CO_2_ before being analysed on a Hydra 20-22 Continuous Flow Isotope Ratio Mass Spectrometer (CF-IRMS; Sercon Ltd., UK).

### Comparative metabolomics

Sealed cultures of *M. smegmatis* were harvested during exponential phase (EXP; O_2_ = 13%, OD_600_ = ∼1.0), the hypoxic transition (TR; O_2_ = 0.5%, OD_600_ = 2.6, 12 hours post OD_max_) and hypoxic stationary phase (ST; O_2_ = 0%, OD_600_ = 2.2) in quadruplicate. 15 mL of culture were pelleted via centrifugation (4,500 ×*g*, 15 min, 4°C) and washed once via resuspension in 15 mL of 1X PBS, pelleting via centrifugation as before, discarding supernatant and flash freezing the pellet in liquid N_2_. Pellets were stored at -80C. Pellets were resuspend thoroughly in 200 µL extraction solvent (2:6:1 chloroform:methanol:water v/v/v and spiked with 2 µM generic internal standard (CHAPS/CAPS/PIPES and Tris) at 4 °C, and subject to three freeze-thaw cycles by snap freezing in liquid N_2_ and then thawing on ice. Samples were mixed on a vibrating mixer (PCV-3000, 1500 RMP, 1 sec, Vortex = Hard, 10, cycle = 60, stop) at 4 °C and then centrifuged at 20,000 *×g*, 10 min, 4 °C). 180 µL of supernatant was transferred into new 1.5 mL tube and frozen at -80 °C. Immediately prior to LC-MS analysis samples were thawed on ice and centrifuged at 20,000 *×g*, before being transferred to sample vial inserts.

LC-MS analysis was performed using a Dionex RSLC3000 UHPLC coupled to a Q-Exactive Plus Orbitrap MS (Thermo). Samples were analysed by hydrophilic interaction liquid chromatography (HILIC) following a previously published method^48^. The chromatography utilized a ZIC-p(HILIC) column 5 µm 150 x 4.6 mm with a 20 x 2.1 mm ZIC-pHILIC guard column (both Merck Millipore, Australia) (25 °C). A gradient elution of 20 mM ammonium carbonate (A) and acetonitrile (B) (linear gradient time-%B: 0 min-80%, 15 min-50%, 18 min-5%, 21 min-5%, 24 min-80%, 32 min-80%) was utilised. Flow rate was maintained at 300 μL/min. Samples were kept in the autosampler (6 °C) and 10µL was injected for analysis. MS was performed at 70,000 resolution operating in rapid switching positive (4 kV) and negative (−3.5 kV) mode electrospray ionization (capillary temperature 3000°C; sheath gas flow rate 50; auxiliary gas flow rate 20; sweep gas 2; probe temp 120 °C). For accurate metabolite identification, a standard library of ∼500 metabolites were analysed before sample testing and accurate retention time for each standard was recorded. This standard library also forms the basis of a retention time prediction model used to provide putative identification of metabolites not contained within the standard library^49^. Acquired LC-MS/MS data was processed in an untargeted fashion using open source software IDEOM, which initially used *msConvert* (*ProteoWizard*)^50^ to convert raw LC-MS files to *mzXML* format and *XCMS* to pick peaks to convert to .*peakML* files^51^. *Mzmatch* was subsequently used for sample alignment and filtering^52^. IDEOM was utilised for further data pre-processing, organisation and quality evaluation^51^. Data was further analysed using MetaboAnalyst^53^ (www.metaboanalyst.ca/): no data filtering was performed (<5000 features). Samples were normalised by median ion peak ion intensity, subject log_10_ transformation and Pareto scaling. Volcano plots for each comparison were generated (fold change threshold = 2.0, p-value (FDR) threshold: 0.01) and .csv were further analysed in Microsoft Excel.

Map annotations from IDEOMv21 were used to broadly group metabolites by predicted function. For more specific groupings (PPP, EDP, glycolysis, TCA), KEGG pathway annotations (https://www.genome.jp/kegg/pathway.html) were used. Data visualisation was performed in GraphPad Prism 9. Spreadsheets used for the processing and analysis of metabolomics data are available in the supplementary.

### Comparative proteomics

Sealed cultures of *M. smegmatis* were harvested in tandem with cultures harvested for comparative metabolomics. 15 mL of culture were pelleted via centrifugation (4,500 x *g*, 15 min, 4°C) and washed three times via resuspension in 15 mL of 1X PBS, pelleting via centrifugation as before, and discarding the supernatant. After the final wash, pellets were resuspended in 1 mL PBS and transferred to a 1.5 mL tube, pelleted via centrifugation (21,000 *×g*, 10 min, 4 °C) before discarding the supernatant and flash freezing the pellet in liquid N_2_.

Cell pellets were solubilised in 5% SDS 100 mM Tris-HCl with heating at 95°C for 10 min to denature enzymes. Subsequently, DNA was sheared using probe sonication, then protein concentration was determined using the Pierce™ BCA Protein Assay Kit as per manufacturer’s instructions (cat:23225, Thermo Scientific™). Samples were then processed using the S-trap protocol as per the manufacturer’s instructions^54^. Eluted peptides were acidified to 1% tri-flouro acetic acid and purified using Stage-tips packed with SDB-RPS (Empore)^55^, iRT peptides were spiked into all samples before LC-MS/MS analysis.

LC MS/MS data was acquired using a Dionex UltiMate 3000 RSLCnano for peptide separation and analysed with an Orbitrap Eclipse Tribrid mass spectrometer (Thermo Scientific) with an Acclaim PepMap RSLC analytical column (75 µm x 50 cm, nanoViper, C18, 2 µm, 100Å; Thermo Scientific) and an Acclaim PepMap 100 trap column (100 µm x 2 cm, nanoViper, C18, 5 µm, 100Å; Thermo Scientific) using a 120 minute linear gradient for separation with two FAIMS compensation voltages (-45, -65)^56^. This was performed by the Monash Proteomics and Metabolomics Facility.

The raw data files were analysed with the Fragpipe software suite 18.0 (MSFragger version 3.5, Philosopher version 4.1.1)^57,58^. The standard label free quantification match-between-runs (LFQ-MBR) workflow was applied with no changes to workflow, employing IonQuant^59^ and the MaxLFQ method of protein abundance calculations^60^. Searches were performed against the *M. smegmatis* proteome database (September 2022) and with common contaminants. The proteomics data were further analysed using LFQ-Analyst which performed data manipulation and statistical tests with standard parameters^61^. Contaminant proteins, reverse sequences and proteins identified “only by site” were filtered out. LFQ was converted to log_2_ scale and missing values were imted using the Missing not At Random (MNAR) method. Protein-wise linear models combined with empirical Bayesian statistics were used for differential expression analyses. The limma package from R Bioconductor was used to generate a list of differentially expressed proteins for each pair-wise comparison. A fold-change threshold of 2 was used and an adjusted p-value threshold of 0.05 (Benjamini-Hochberg method) was used to identify differentially abundant proteins^62^.

UniProt accession IDs were used to match *M. smegmatis* gene numbers (MSMEG_XXXX) with KEGG Ortholog (KO) identifiers and KEGG module numbers. KEGG module numbers were used to annotate *M. smegmatis* genes with associated KEGG pathways. Spreadsheets used for the processing and analysis of proteomics data are available in the supplementary. The mass spectrometry proteomics data have been deposited to the ProteomeXchange Consortium via the PRIDE^63^ partner repository with the dataset identifier PXD045129.

### Electron cryotomography data collection and processing

Sealed cultures of *M. smegmatis* were harvested during mid-exponential phase (EXP; O_2_ = 17%, OD_600_ = 0.5), the hypoxic transition (TR; O_2_ = 0.7 %, OD_600_ = 2.5) and mid-stationary phase (ST; O_2_ = 0%, OD_600_ = 1.9, 5 days post OD_max_). About 4 µl of aerated and hypoxia samples were pipetted on glow-discharged Quantifoil Au extra thick EM-grids (R2/2, 200 mesh, Electron Microscopy Sciences). The extra fluid was blotted (blot force of 7 s and blot time 5 sec) using Whatman filter paper and plunge-frozen in liquid ethane using a FEI Vitrobot Mark IV, set to 22 °C and 100% humidity.

Data were collected using a 300 kV FEI Titan Krios TEM equipped with a Gatan K3 direct detector and Gatan energy filter (slit width of 20 kV). Tilt-series were collected in movie mode with a total electron dose of 150 e−/Å^2^, defocus of -8 µm, and pixel size of 3.4 Å.

Motioncor was used for motion correction^64^. Tilt series were aligned with IMOD and subsequent 2K binned micrographs were reconstructed using Tomo 3D^65,66^. Tomograms were segmented using Dragonfly ((Object Research Systems (ORS) Inc, Montreal, Canada, software available at www.theobjects.com/dragonfly), an U-Net convolutional neural network-based software. Tomograms were loaded into Dragonfly and filtered using the histogram equalization filter followed by a 3D Gaussian filter to boost contrast. The feature of interest was initially hand segmented to generate a multi ROI training output. This was then used for unsupervised and unbiased segmentation.

## Acknowledgements

This work was supported by three National Health & Medical Research Council Emerging Leader Fellowships (APP1178715 to C.G.; APP1197376 to R.G. and APP1196924 to D.G.), three Australian Research Council Discovery Project grants (DP200103074 to C.G. and R.G.; DP230103080 to C.G. and R.G.; DP210101595 to C.G. and P.L.M.C.), and an Australian Government Research Training Stipend Scholarship (to D.L.G.). We thank Matthew Johnson and Ian Holmes Imaging Center (Bio21 Institute, University of Melbourne) for help with cryo-ET data collection.

## Author contributions

C.G. conceived and oversaw the study. C.G., D.L.G., P.L.M.C., R.G., and D.G. designed experiments. D.L.G. and J.L. was responsible for growth, oxygen consumption, and hydrogen production experiments. D.L.G., T.H., J.L., W.W.W., and P.L.M.C. conducted isotopologue analysis. D.L.G., H.L., J.R.S., I.H., E.T., R.B.S., and C.B. conducted proteomic and metabolomic analysis. M.M, D.G., and D.L.G. conducted cryo-ET analysis. T.D.W. and L.J. conducted lipid analysis. D.L.G. and C.G. wrote the manuscript with input from all authors.

## Supplementary Note 1: Additional proteomic and metabolomic analyses

The proteomic response of *M. smegmatis* differs in fundamental ways to *M. tuberculosis*. Studies in *M. tuberculosis* have demonstrated that rerouting of the carbon flux from the TCA cycle to the glyoxylate shunt and methylitrate pathways are key parts of the pathogen’s response to oxygen depletion, and contrary to our findings, isocitrate lyase (*icl*) is upregulated during hypoxia and essential during infection^14,16,67–69^. It has also been shown these pathways drive an accumulation of succinate during oxygen depletion, which *M. tuberculosis* excretes to maintain membrane potential^14,67^. Consistent with the downregulation of *icl* during oxygen depletion in *M. smegmatis*, we observed no accumulation of succinate, and in fact detected a 60-fold reduction of succinate in the stationary phase vs exponential phase comparison (**Fig. 3**). These differences may be in part due to the fatty acid rich diet of *M. tuberculosis* during infection, as the methylcitrate pathway is required for the catabolism of odd-chain fatty acids^68,70^, and the glyoxylate shunt is required for the regeneration of TCA cycle intermediates when relying on fatty acids for energy and carbon^67^. The differences may also be due to additional metabolic flexibility conferred by H_2_ fermentation, as *M. tuberculosis* excretes succinate during hypoxia in order to maintain membrane potential and ATP production^14,67^, while *M. smegmatis* can continually produce ATP via substrate level phosphorylation by dispensing of reducing equivalents via Hyh. With respect to the DosR-regulated genes, MSMEG_3934 is annotated as a phosphoenolpyruvate synthase (PEPS) that catalyses the gluconeogenic reaction ATP + H_2_O + pyruvate → AMP + 2H^+^ + phosphate + phosphoenolpyruvate^71,72^. PEPS has no homolog in *M. tuberculosis,* and its predicted function has not been experimentally investigated, and considering it is the second most upregulated protein in the hypoxic transition vs exponential phase comparison, it is somewhat difficult to reconcile with our data where we observe a decrease in abundance in phosphoenolpyruvate (3-fold and 18-fold, during hypoxic transition vs exponential phase and stationary phase vs exponential phase, respectively), while observing no changes to the abundance of pyruvate (**Fig. 3**). Although we observed a decrease in the abundance of PEP during oxygen depletion, and no change in the abundance of pyruvate, PEPS mediated flux may happen very early in the response to hypoxia to arrest growth. Similarly, in *M. tuberculosis,* which lacks a PEPS homolog, the metabolic rerouting towards TAG synthesis plays a key role in growth arrest by depleting PEP^73^ and acetyl-CoA^74^, and rerouting towards the PPP is mediated by pyruvate kinase^41^. We can only speculate about possible posttranscriptional regulation when the abundance of metabolites and associated enzymes do not correlate, though investigation of these mechanisms require intensive biochemical characterisation^75^.

The homolog of PfkB in *M. tuberculosis* is also upregulated during hypoxia, can catalyse the reverse gluconeogenic reaction (albeit with low efficiency), and exhibits lower activity, sensitivity to allosteric regulators and substrate specificity than PfkA, which is essential for growth on glucose^75,76^. We did not identify PfkA in our proteomics dataset, but it was not found to be differentially regulated in other microarray^34^ and transcriptomic^77^ datasets comparing hypoxic and aerated cultures of *M. smegmatis*. The glycolysis enzyme PksB potentially has an allosteric role given its lower activity, ability to catalyse the reverse gluconeogenic reaction, and recalcitrance to metabolites in *M. tuberculosis*. It may allosterically regulate its isozyme PksA, playing a pivotal role in redirecting glycolytic flux to the PPP / EDP, if these properties are conserved in *M. smegmatis*.

We also observed changes in cofactor levels during hypoxia based on the metabolomics. Both the PPP and EDP reduce NADP^+^ to NADPH, which in turn provides reductive power for anabolism. During the hypoxic transition vs exponential phase comparison, we only observed changes to NAD^+^/NADH, which both increased in abundance by 2-fold (**Fig. 3**). However, we observed a dramatic decrease in the abundance of both NADP^+^/NADPH (80- to 92-fold respectively), in addition to NAD^+^/NADH (38- to 13-fold, respectively) and ADP (9-fold, ATP not detected), during the stationary phase vs exponential phase comparison (**Fig. 3**). This likely reflects a dramatic downshift in overall metabolism and energy expenditure as *M. smegmatis* enters non-replicative persistence. Due to the nature of our comparative analysis, compared to absolute quantification of metabolites, we were unable to calculate the stoichiometric ratios of each redox cofactor pair.

